# An intra-oral flavor discrimination task in freely moving mice

**DOI:** 10.1101/2023.05.26.542382

**Authors:** Kazuki Shiotani, Yuta Tanisumi, Yuma Osako, Koshi Murata, Junya Hirokawa, Yoshio Sakurai, Hiroyuki Manabe

## Abstract

Flavor plays a critical role in the pleasure of food. Flavor research has mainly focused on human subjects and revealed that many brain regions are involved in flavor perception. However, animal models for elucidating the mechanisms of neural circuits are lacking. Herein, we demonstrate the use of a novel behavioral task in which mice are capable of flavor discrimination. When the olfactory pathways of the mice were blocked, they could not perform the task. However, behavioral accuracy was not affected when the gustatory pathway was blocked by benzocaine. These results indicate that the mice performed this discrimination task mainly based on the intraoral flavor. We conclude that this novel task can contribute to research on the neural mechanisms of flavor perception.

## Introduction

Flavor perception is an important factor in determining the pleasure of eating, which is the most complex and active behavior in communication with the outside world. Almost all senses are involved in this perceptual process, but olfaction is the most important one^1^. However, the dominant role of olfaction in flavor perception during natural interactions with the environment has not yet been fully evaluated. In humans, odor molecules generated in the mouth have been shown to have a significant effect on food perception^2–4^, and this effect occurs only during the active state of breathing out through the nose during mastication and swallowing^5^. Many studies have identified sub-areas where neural activity is driven by food flavors with active sensing, such as the olfactory cortex (OC), anterior cingulate cortex, orbitofrontal cortex, and insular cortex^1, 6–9^. However, there are limitations in using humans to develop neural circuit studies that focus on olfaction in flavor perception. This is because human neuroimaging studies do not allow for invasive experimental methods, the spatiotemporal resolution of measuring neural activity is low, and verification in the environment, such as pharmacological or physical impairment of odor or taste perception, where the flavor experience is properly controlled are difficult.

To overcome these problems, we demonstrated that mice are an effective animal model for flavor research in natural environments. Previous studies have reported indirect evidence that rodents perceive odor molecules generated in their mouths^10–14^. Steady airflow models of the nasal cavity also support the relevance of intraoral olfactory perception, although with less efficiency than that in humans^15, 16^. Although flavored odor discrimination tasks using rodents have been proposed^17–19^, there are still some limitations. First, a fixed-head system limits active behavior in the natural environment, which is crucial for flavor perception. Therefore, it is necessary to design a behavioral paradigm in which animals can freely access flavor stimuli and then make a decision when they have completed intraoral olfactory perception in sucrose water of their own will. Second, the previous task protocols did not verify whether rodents specifically exhibited negative effects on flavor perception when their sense of olfaction was impaired. They performed bilateral olfactory bulb (OB) removal to demonstrate the dominance of olfaction in flavor perception; however, they could not reduce or evaluate the effects of the rodents’ physical condition associated with olfactory loss because the removal of bilateral OBs in rodents is known to be a depressive model^20, 21^. Third, by dissolving odor molecules in water, they were unable to fully rule out the possibility that rodents discriminate based on taste or somatosensory perception.

To address these problems, we established a new behavioral task for freely moving mice and verified whether they could discriminate between flavored and non-flavored. After confirming that the mice were capable of performing the task, pharmacological and lesion experiments were conducted. When the olfactory system was blocked with an olfactotoxic drug or with unilateral OB removal, the percentage of correct responses decreased significantly. When the gustatory system was blocked with a topical sodium channel blocker, the percentage of correct responses decreased slightly. These results indicate that the mice performed this task primarily using flavor perception, consisting of a retronasal olfactory sensation.

## Results

### Flavor discrimination task

We conducted a behavioral experiment in which motivated animals (thirsty) were trained to detect flavored sucrose water (sucrose water dissolved in an odorant) and non-flavored sucrose water. Turning on a light on the right stimulus port instructed the mouse to start the trial, approach the stimulus port, and poke the nose into the stimulus port (Figure 1A). To prevent the mouse from using odor wafting from the flavored water as a reference for their choice, the port was filled with odor dissolved in the flavored water immediately before presenting the test stimulus. The concentration of the odor presented by the olfactometer was 100 times higher than the concentration of the odor dissolved in the flavored sucrose water. An average of 350 ms after odor delivery, either the flavored or non-flavored sucrose water was presented. One second after the onset of flavor stimulation, the light was turned off and the mouse withdrew its nose from the stimulus port. If the go cue stimulus (flavored sucrose water, conditioned stimulus +: CS+) was presented, the mouse was required to move and poke its head into the left reward port within 5 s to obtain sucrose water as a reward (Figures 1A and B). At the reward port, the mouse was required to maintain its head in the port for 500 ms while waiting for the sucrose water delivery. A drop of sucrose water was administered as a reward 500 ms after nose poking. If a no-go cue stimulus (non-flavored sucrose water, CS-) was presented, the mouse was required to prohibit poking its head into the water-reward port for 5 s after the end of non-flavored sucrose water delivery. Response time was defined as the time between the CS onset and withdrawal of the nose from the stimulus port, and moving time was defined as the time between the withdrawal of the nose from the stimulus port and poking of the reward port (Figure 1A). In addition, when CS+ was presented, the condition in which the mouse performed go behavior was defined as Hit, and the condition in which the mouse exhibited no-go behavior was defined as Miss. When CS-was presented, the condition in which the mouse exhibited no-go behavior was designated as Correct Rejection (CR), and the condition in which the mouse performed go behavior was designated as False Alarm (FA) (Figure 1B).

**Figure. 1.**
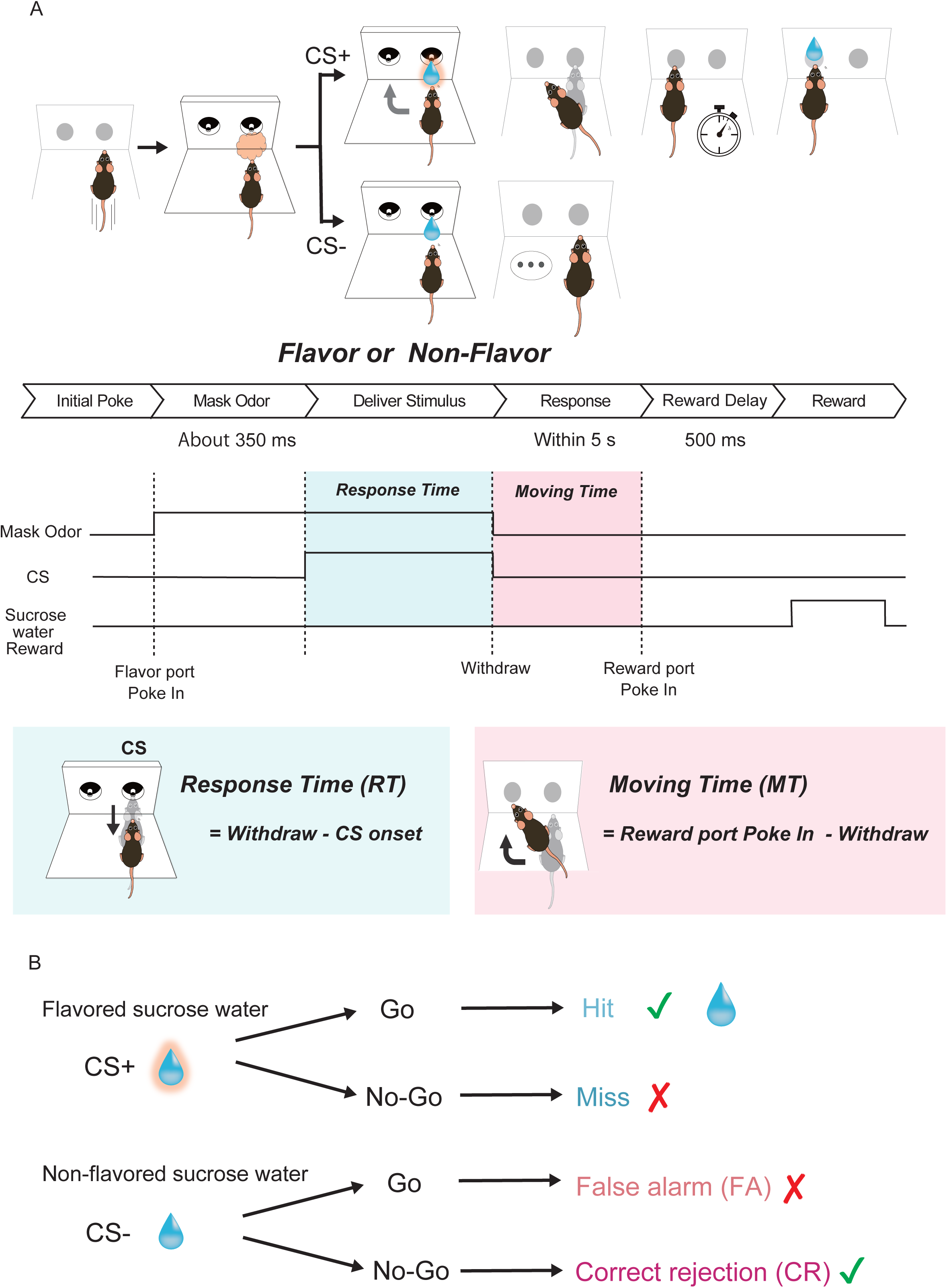
Schematics of sequences of behavioral events in the flavor discrimination task. (**A**) Time course of the flavor discrimination task. Behavioral epoch temporal progression from left to right. The mice initiated the trial by poking their nose into the stimulus port and were presented the masking odor (the same odor dissolved in the water) for about 350 ms. After that, flavored sucrose water or sucrose water (6 μl) was delivered in the same port. If flavored sucrose water (go cue water) was presented, the mice were required to move to and poke their nose into the left water reward port within 5 s. At the reward port, the mice were required to keep poking their nose for 500 ms before water delivery began. Then 12 μl of water was delivered as a reward. If sucrose water (no-go cue water) was presented, the mice were required to avoid entering the reward port for 5 s. (**B**) Trial outcomes in go and no-go trials.

### Behavioral accuracy and response time in flavor discrimination task

Behavioral accuracy (% correct response) steadily increased throughout the training (an example mouse A: Figure 2A). The behavioral accuracy of the last training session was significantly higher than that of the first training session in all mice (n = 40 mice, Figure 2B, ***p < 0.001, paired t-test). Approximately 21 sessions were required to complete the learning task (>75% correct). As response time decreases as learning progresses^22^, we examined response time as an indicator of learning progress and task difficulty. The response time in the last training session was significantly lower than that in the first training session (Figure 2C, ***p < 0.001, Wilcoxon rank-sum test). In the last training session, when the mice were well-trained, the behavioral accuracy remained above 75% in each block (20 trials per block) within a session (average of 40 mice, Figure 2D).

**Figure. 2.**
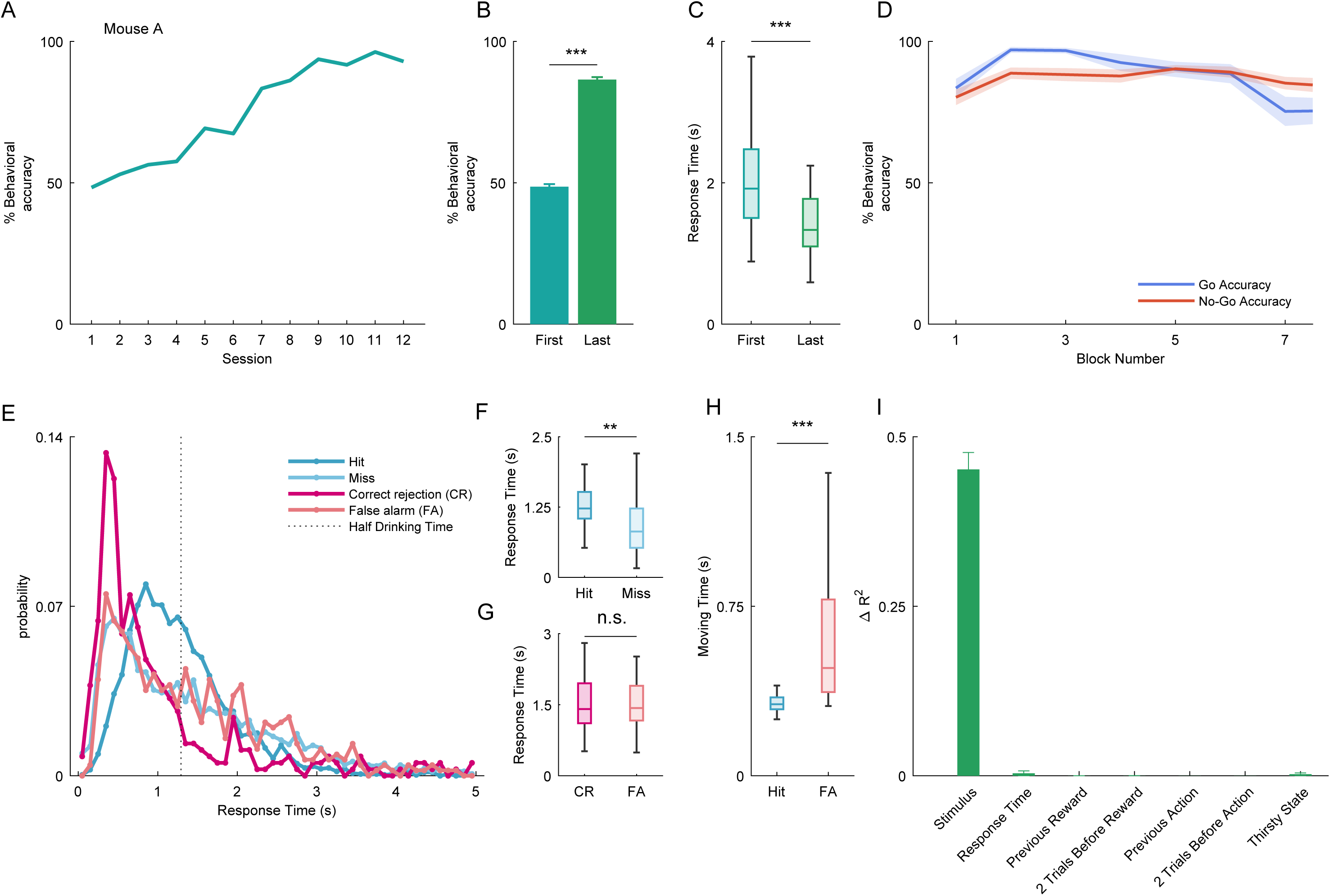
Behavioral performance in the flavor discrimination task. (A) Performance of an example mouse A in the flavor discrimination task. (B) Average performance in the first and last training session (n = 40 mice, ***p < 0.001, paired t-test). Error bars show SEM. (C) The average response time (from the onset of stimulus presentation to the timing of nose withdrawal) in the first and last training session (***p < 0.001, Wilcoxon rank sum test). (D) Time series of the average block (one block: 20 trials) performance in the last training session (n = 40 mice). The average accuracies are shown (blue line, go action; red line, no-go action). Error shades are shown SEM. (E) Histogram of the response time in the last session (blue line, Hit; light blue line, Miss; red line, correct rejection (CR); light red line, false alarm (FA)). The dotted line indicates the half of average drinking time (see STAR Methods). (F) The boxplot of response times for Hit and Miss in the last training session (**p < 0.01, Wilcoxon rank sum test). (G) The boxplot of response times for CR and FA in the last training session (n.s., not significant, Wilcoxon rank sum test). (H) The boxplot of moving time for Hit and FA in the last training session (***p < 0.001, Wilcoxon rank sum test). (I) Impact of task parameters on behavioral variability (see STAR Methods) using a GLM fitting for go or no-go actions. Each average Δ*R*^2^ is shown. Error bars show SEM.

To investigate the key components of task performance in this behavioral task, from stimulus to action, we examined the response time in the last training session (Figure 2E). The vertical dashed line indicates the average duration when the mice drank 6 µl of sucrose water as a reward, calculated as half of the time from the onset of drinking the 12 µl sucrose reward after stimulus presentation at the reward port exit (n = 40 mice). Note that the duration of drinking flavored sucrose water as a stimulus to predict go/no-go behavior was shorter than that of drinking sucrose water as a reward, suggesting that the mice understood the context in relation to how each water was used and were in a behavioral state during stimulus sampling, including not only drinking sucrose water but also perceptual detection of the flavored stimulus. The response time in the go action was different between Hit and Miss, whereas the response time in the no-go action was not different between CR and FA (Figure 2F and G; **p < 0.01 and not significant, respectively; Wilcoxon rank-sum test). There was no difference between the correct condition (Hit and CR combined) and false condition (Miss and FA combined) (Figure S1B, not significant, Wilcoxon rank-sum test). The moving time in the Hit cases was significantly shorter than that in the FA cases (Figure 2H, ***p < 0.001, Wilcoxon rank-sum test). These results indicate that the mice were able to take time to taste and quickly move on to the next behavior when they correctly recognized the flavored sucrose water.

To determine what drove the mice’s choice, we applied a generalized linear model (GLM) analysis to the behavioral data. We used multiple variables, such as current stimulus, response time, previous reward, two trials before reward, previous action, two trials before action, and thirsty state as independent variables and predicted go or no-go responses to flavored and non-flavored stimuli (Figures 2I and S1C; see STAR Methods). We compared the variance uniquely explained by the variable (see STAR Methods). Note that the individual delta-R^2^ does not add to the total R^2^ because some of the variance can be explained by multiple factors. Most of the action variance was accounted for by the stimulus alone (45.2 %), with reaction time contributing 0.364% and thirsty state contributing 0.226 % (Figure 2I). Together, the responses of the mice in the flavor task were predominantly driven by flavor stimulus information, with little influence from the action history (e.g., previous reward and previous action) and state (e.g., thirsty state).

However, there is a possibility that the mice might distinguish the stimulus by other sensory modalities (e.g., taste and somatosensory). Did they discriminate between stimuli based on flavor differences? We investigated whether olfaction was critical for flavor discrimination by blocking the olfactory pathway in two ways.

### Injection of methimazole to prevent distinguishing between the flavors

Methimazole was used to pharmacologically block the olfactory system. Methimazole, an olfactotoxic drug, disrupts the existing olfactory sensory neurons (OSNs) almost evenly throughout the olfactory epithelium (OE) by activating the apoptotic cascade in OSNs^23^. Since progenitor basal cells in the OE remained intact after 3 days of methimazole-induced injury, they produced newly generated OSNs, and the OE returned to its pre-injury level in approximately 28 days^24^. After the mice successfully learned this behavioral task, one group of mice was injected with methimazole, and another group of mice was injected with saline as a control (Figure 3A). The sessions before, one day after, two days after, and three days after drug administration were called the pre, post1, post2, and post3 sessions, respectively. In the methimazole-treated group, the percentage of correct answers decreased post1 session, and it decreased significantly in the post2 and post3 sessions compared to the control group (Figure 3B; **p<0.01 and ***p<0.001, respectively; Bonferroni’s multiple comparison test). To examine changes in the behavioral responses of the mice to methimazole administration, the number and percentage of the conditions were calculated pre and post3 sessions in the methimazole administration group. We found that the number of Hit significantly decreased and the number of Miss increased in the pre and post3 sessions (Figures 3C and D, *p<0.05, paired t-test). In addition, mice in the control group were intraperitoneally administered methimazole after data were obtained from the saline administration experiment. Behavioral accuracy significantly decreased at post1, post2, and post3 sessions after methimazole administration (Figure S2A; **p<0.01 and ***p<0.001, respectively; Tukey’s test). Similar results were obtained in the control group (Figure S2B and C). These results indicate that both groups were unable to discriminate flavor after pharmacologically blocking the olfactory system.

**Figure. 3.**
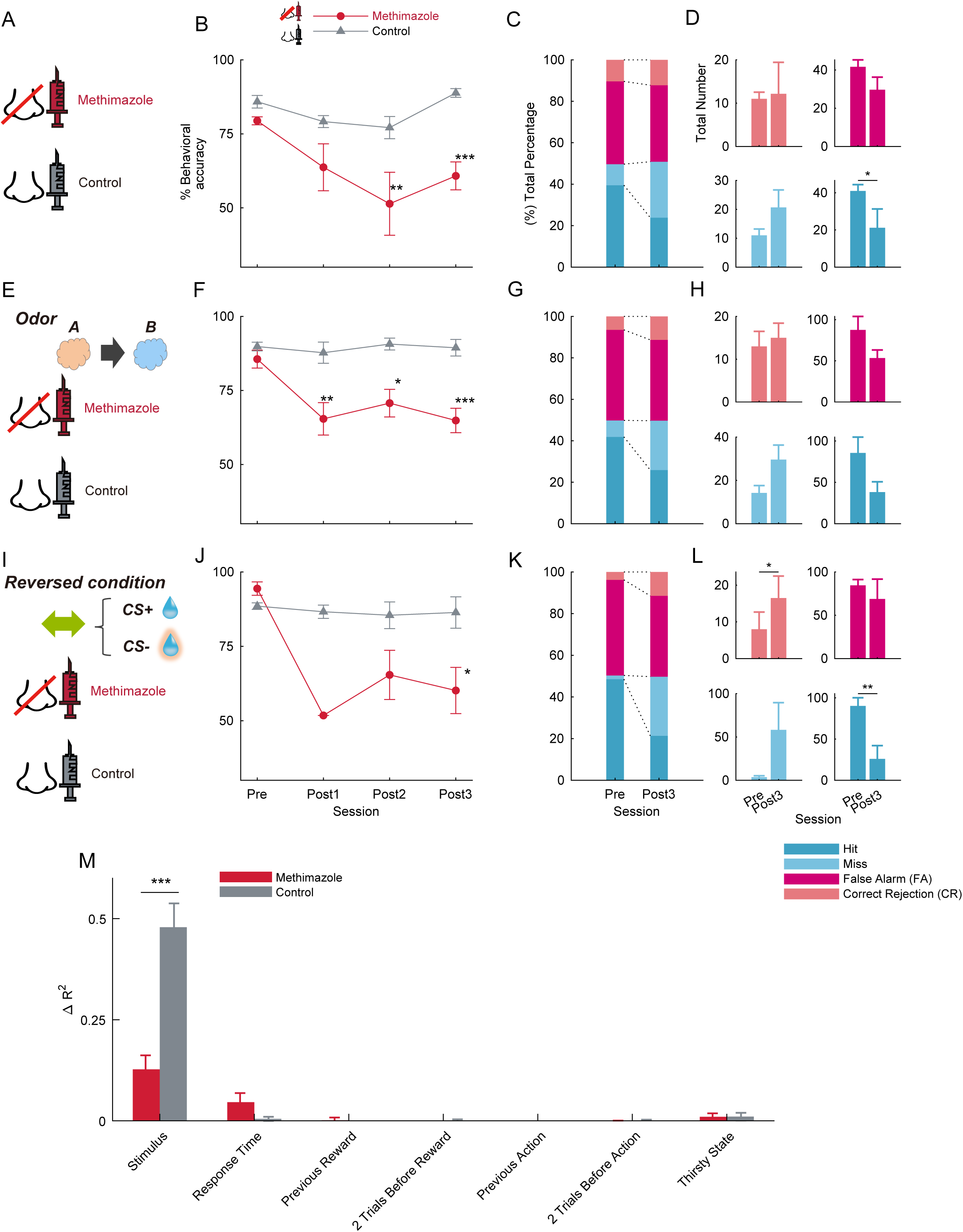
Effects of intraperitoneal methimazole, a disruptor of olfactory cells, on the flavor discrimination task. (A) Schematic of the methimazole (red, n = 6 mice) and saline intraperitoneal injection (gray, n = 6 mice) group. Intraperitoneal injection was done after the pre session was finished. (B) The time course of average behavioral accuracy in the methimazole (red circle line) and control (gray triangle line) groups. The behavioral accuracy in the methimazole group was significantly decreased than that of the control group in the post2 and post3 sessions (*p<0.05; **p<0.01; ***p<0.001, Bonferroni’s multiple comparison test). Error bars show SEM. (C) Total percentage bar showing the proportion of the behavioral action in pre and post3 sessions of the methimazole group (blue, Hit; light blue, Miss; red, correct rejection (CR); light red, false alarm (FA)). (D) The average of the total number bar showing the actual number of trials in which behavioral actions were taken and corresponding to the total percentage bar (Figure 3C) (*p<0.05; **p<0.01, paired t-test). Error bars show SEM. (E-H) Same as in (A-D) but with eugenol as the odor (methimazole group n = 5 mice, control group n = 5 mice). (I-L) Same as in (A-D) but with the opposite relationship between go and no-go stimuli (methimazole group n = 4 mice, control group n = 4 mice). (M) Effect of task parameters on behavioral variability with methimazole and saline administration using a GLM fitting for go or no-go actions (see STAR Methods). Each average Δ*R*^2^ in the post3 session (red, methimazole group n = 13 mice; gray, control group n= 15 mice) is shown (*p < 0.05; **p < 0.01; ***p < 0.001, two-sample t-test). Error bars show SEM.

It is possible that mice can discriminate flavor stimuli based on specific odorants (odor A: amyl acetate). To exclude this possibility, we used another odorant (odor B: eugenol) (Figure 3E). In the flavor task, the external odor stimulus was the same odor used to dissolve the odorant in the flavored sucrose water. Similarly, in this odorant condition, behavioral accuracy at post1, post2, and post3 sessions in the methimazole group was significantly lower than that in the control group (Figure 3F; *p<0.05, **p<0.01, and ***p<0.001, respectively; Bonferroni’s multiple comparison test). After methimazole administration, the percentage of Hit tended to decrease and the percentage of Miss tended to increase in the pre and post3 sessions (Figure 3G and H).

To confirm whether the mice could learn the opposite relationship between go and no-go stimuli in this task, and to verify the effect of methimazole administration in the opposite condition, the CS condition was swapped from the beginning of the training in the flavor task (Figure 3I). Note that we did not conduct reversal learning, in which the CS condition was reversed during the behavioral task. Even when the CS was swapped, the mice were able to learn on the task and methimazole administration showed similar results (Figure 3J, K, and L).

We analyzed whether there was a difference in the response time of the methimazole group before and after administration (Figure S2D, E, F, and G). In the post3 session of the methimazole group, the response times of all responses (Hit, Miss, CR, and FA) were significantly increased compared to those in pre session (Figure S2F; **p < 0.01 and ***p < 0.001, respectively; Wilcoxon rank-sum test), and the moving times of Hit and FA in the post3 session were also significantly longer than those in pre session (Figure S2G; **p < 0.01 and ***p < 0.001, respectively; Wilcoxon rank-sum test). The increase in response and moving time in the post3 session in the methimazole group may reflect the increased difficulty of the behavioral task due to olfactory deprivation.

To determine whether the behavior of mice after methimazole administration was dependent on the stimulus, we applied a GLM analysis to the behavioral data. We used the same multiple variables as described above (Figure 2I and S1C) and predicted go or no-go responses to flavored and non-flavored stimuli (see STAR Methods). To evaluate the effect of methimazole administration on behavior, the stimulus component in the action variance of the methimazole group was significantly decreased compared to that of the control group, and other components, such as the response time in the action variance, were increased (Figure 3M, ***p < 0.001, two-sample t-test). Taken together, blocking the olfactory pathway with methimazole rendered mice unable to discriminate between the presence and absence of flavors. These results indicated that the mice discriminated flavors through their olfactory system.

### Olfactory sensory neurons regenerate after 28 days, recovering to pre-methimazole performance

Newly generated OSNs compensate for methimazole-induced loss of OSNs within 28 days^24^. We examined the performance accuracy approximately 28 days after methimazole administration to verify whether it had recovered compared to that at 3 days after the injection (see STAR Methods). We compared the behavioral accuracy before methimazole administration (pre), 3 days after methimazole-induced injury (post3), and 28 days after methimazole-induced injury (post28). Surprisingly, the behavioral accuracy of the post28 session increased significantly compared to that of the post3 session and did not differ from that of the pre session (Figure 4A; **p<0.01 and ***p < 0.001, respectively; Tukey’s test). This result showed that the behavioral accuracy after 28 days of methimazole treatment returned to that before methimazole treatment. We examined the behavioral responses of the mice to determine whether there were any differences between the pre and post28 sessions. The individual percentage of actions taken during the behavioral task in the post28 session was different from that in the post3 session but not from that in the pre session (Figure 4B). Furthermore, the response and moving time in the post28 session were also significantly different from those in the post3 session but not significantly different from those in the pre session (Figure 4C and D; *p<0.05 and ***p < 0.001, respectively; Tukey’s test). To clarify what the behavior of the mice was based on in the post28 session and whether there was any difference between the pre and post28 sessions, we applied a GLM analysis to the behavioral data. We demonstrated that the stimulus component of the action variance dropped in the post3 session, but recovered to its pre session state in the post28 session (Figure 4E, *p < 0.05, **p < 0.01, Tukey’s test). These results showed that mice in the post28 session returned to the behavioral conditioning before methimazole administration and could discriminate between flavor differences. These results indicate that even if most OSNs are destroyed and the nerves between the OSNs and olfactory bulb (OB) are rewired, it is possible to take appropriate actions based on the flavor.

**Figure. 4.**
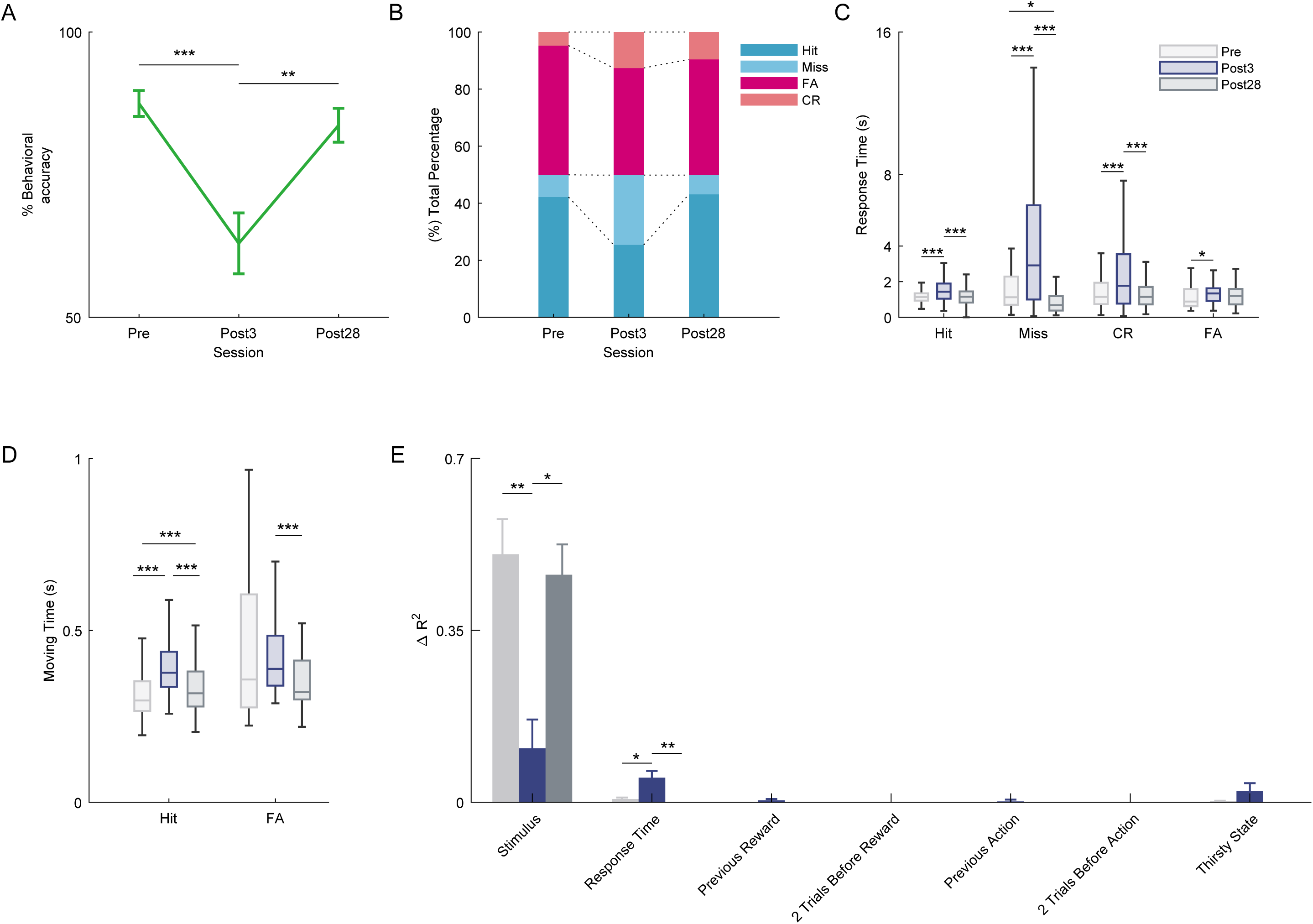
Recovery of behavioral performance to pre-injection performance after 28 days of methimazole injection. (A) Average performance in the pre, post3 and post28 days in the methimazole group (n =6 mice, **p<0.01, ***p < 0.001, Tukey’s test). Post28 shows the 28th day after the day of methimazole injection. Error bars show SEM. (B) Percentage bar showing the percentage of the behavioral action in pre, post3 and post28 days (blue, Hit; light blue, Miss; red, correct rejection [CR]; light red, false alarm [FA]). (C) The boxplot of response time for individual actions during pre (gray), post3 (dark blue), and post28 (dark gray) days (*p<0.05, ***p < 0.001, Tukey’s test). (D) The boxplot of moving time for go actions (Hit and FA) during pre (gray), post3 (dark blue), and post28 (dark gray) days (***p < 0.001, Tukey’s test). (E) Effect of task parameters on behavioral variability over time on the day of methimazole administration is shown using a GLM fitting for go or no-go actions. Each average Δ*R*^2^ (gray, pre day; dark blue, post3 day; dark gray, post28 day) is shown (*p < 0.05, **p < 0.01, Tukey’s test). Error bars show SEM.

### OB removal group showed a dramatic decrease of performance accuracy in the task

To rule out the possibility that methimazole could affect regions other than the OSNs, we blocked the olfactory pathway using other methods. We adopted an irregular olfactory deprivation method that combined unilateral OB removal and opposite naris occlusion. If the bilateral OB are destroyed, the mortality rate following the operation is higher, and the mice are known to suffer from depression^20, 21^. To avoid the effects of depression on the performance, we removed the unilateral OB and inserted a custom-made 10 mm silicon tube into another nostril to block the air flow. A sham operation was performed in the same manner, but the OBs were left intact, as in the control group (Figure 5A). Behavioral training was started three days after surgery to avoid the influence of olfactory complements due to plastic changes in other neural circuits. After behavioral accuracy reached at least 75%, a silicone tube was inserted into the nostril on the contralateral side of the removal site of each mouse using a previously reported procedure^25, 26^ with tweezers. The behavioral accuracy of the OB removal group was significantly lower than that of the control group in the post session (Figure 5B, *p<0.05, Bonferroni’s multiple comparison test). The number of Hit and CR in the post session were significantly lower than those in the pre session (Figure 5C and D, *p<0.05, paired t-test). These results indicate that olfaction is necessary for discrimination in flavor discrimination tasks.

**Figure. 5.**
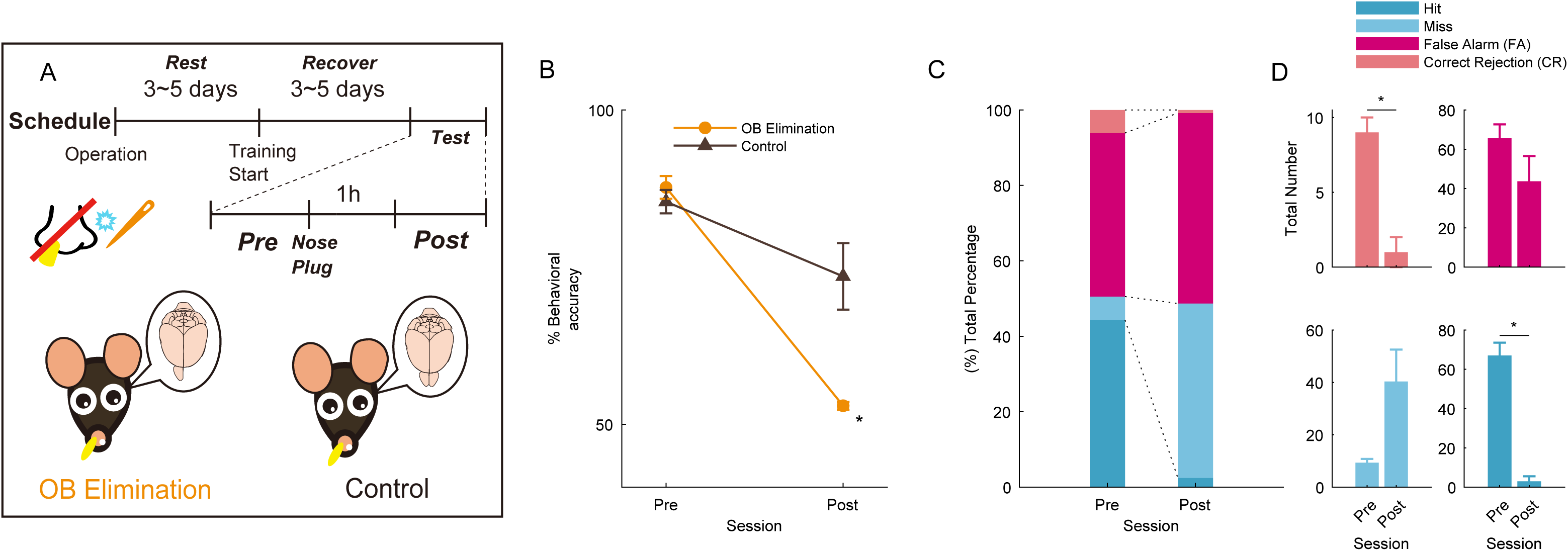
Effects of olfactory bulb elimination on the flavor discrimination task. (A) Schematic of the olfactory bulb (OB) elimination (n = 3 mice) and control (shame operation) groups (n = 7 mice). The test session was divided into two days. The session on the day before the nasal plug was designated as the pre session, and the session 1h after the nasal plug was designated as the post session. (B) The average behavioral accuracy in the OB elimination (orange circle line) and control (gray triangle line) groups before and after the nasal plug procedure. The behavioral accuracy in the OB elimination group was significantly decreased than that in the control group in the post session (*p<0.05, Bonferroni’s multiple comparison test). Error bars show SEM. (C) Total percentage bar showing the proportion of the behavioral action in pre and post session of the OB elimination group (blue, Hit; light blue, Miss; red, correct rejection [CR]; light red, false alarm [FA]). (D) The average of the total number bar showing the actual number of trials in which behavioral actions of the OB elimination group were taken and corresponding to the total percentage bar (*p<0.05, paired t-test). Error bars show SEM.

### Benzocaine group showed a little decrease of performance accuracy in the task

To clarify the effect of dissolved odor molecules on the taste and somatosensory perception of the tongue, 20% benzocaine (a topical sodium channel blocker) was applied to the tongue. On the test days, after the mice were well-trained, benzocaine was applied to the exposed dorsal surface of their tongue immediately after 50 trials (Figure 6A). To prove that the drug works, another group of mice was trained to learn taste discrimination using a go/no-go task and were administered benzocaine on the test days. In the taste-discrimination task, the go cue stimulus was 300 mM sucrose water and the no-go cue stimulus was 154 mM NaCl water^27^ (Figure S3A). The other parameters were the same as those used in the flavor discrimination task (see STAR Methods). In the taste-discrimination task, behavioral performance after benzocaine treatment decreased significantly compared to that before benzocaine treatment (Figure S3B). However, in the flavor discrimination task, behavioral performance after benzocaine treatment was not significantly different from that before benzocaine treatment (Figure 6B). The behavioral responses of mice after benzocaine administration in the flavor discrimination task did not change as dramatically as that in the taste discrimination task (Figure 6C, 6D, S3C, and S3D). With benzocaine administration, the response time changed, and the moving time in the Hit was prolonged in both behavioral tasks (Figure 6E, 6F, S3E, and S3F). Furthermore, when we examined what the mice discriminated in the pre- and post-sessions by administering benzocaine, we found that the stimulus component of the action variance in the flavor discrimination task did not change significantly in the pre- and post-sessions, while it changed significantly in taste-discrimination task (Figure 6G and S3G, *p < 0.05, two-sample t-test). These results indicate that oral gustatory and somatosensory stimulation by dissolved odor molecules is not a major component in discriminating between the presence and absence of flavors.

**Figure. 6.**
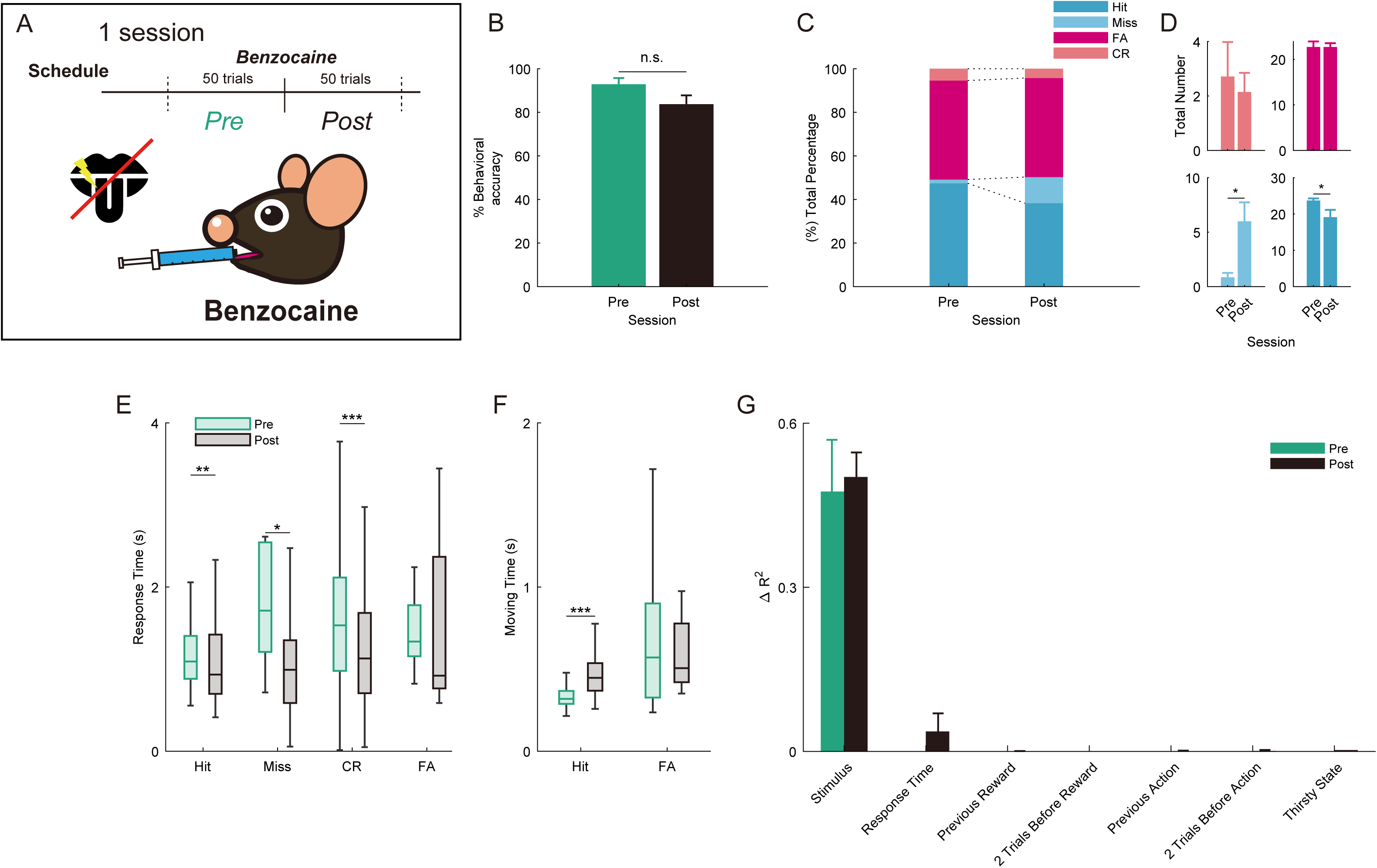
Effects of benzocaine on the flavor discrimination task. (A) Schematic of the schedule of the benzocaine group (n =7 mice). The pre session (50 trials) represents the session before the benzocaine procedure. The post session (50 trials) represents the session after benzocaine was administered to all mice immediately after the pre session in the same day. (B) The average behavioral accuracy in the pre (green) and post (black) sessions of the benzocaine procedure. The behavioral accuracy in the pre session was not significant compared to that in the post session (n.s. not significant, paired t-test). Error bars show SEM. (C) Percentage bar showing the percentage of the behavioral action in the pre and post sessions (blue, Hit; light blue, Miss; red, correct rejection [CR]; light red, false alarm [FA]). (D) The average of the total number bar showing the actual number of trials in which behavioral actions were taken and corresponding to the total percentage bar (Figure 6C) (*p<0.05, paired t-test). Error bars show SEM. (E) The boxplot of response time for individual actions in the pre (green) and post (black) sessions (*p<0.05; **p<0.01; ***p < 0.001, Wilcoxon rank sum test). (F) The boxplot of moving time for Hit and FA in the pre (green) and post (black) sessions (***p < 0.001, Wilcoxon rank sum test). (G) Effect of task parameters on behavioral variability with benzocaine administration using a GLM fitting for go or no-go actions. Each average Δ*R*^2^ (green, pre session; black, post session) is shown (paired t-test). Error bars show SEM.

### Differences in the accuracy to behavioral task by various validation methods

We have shown how mice discriminate between flavor and non-flavored stimuli in this task in various ways. We also calculated the change in accuracy of the flavor discrimination task using various validation methods to verify which factors had a greater influence. The OB elimination + nose plug effect, followed by the methimazole effect, reduced the accuracy (Figure 7). The unilateral nasal plug was found to reduce the accuracy, and benzocaine had the least effect on the accuracy in this task. These results indicated that olfaction is the most important factor in flavor perception.

**Figure. 7.**
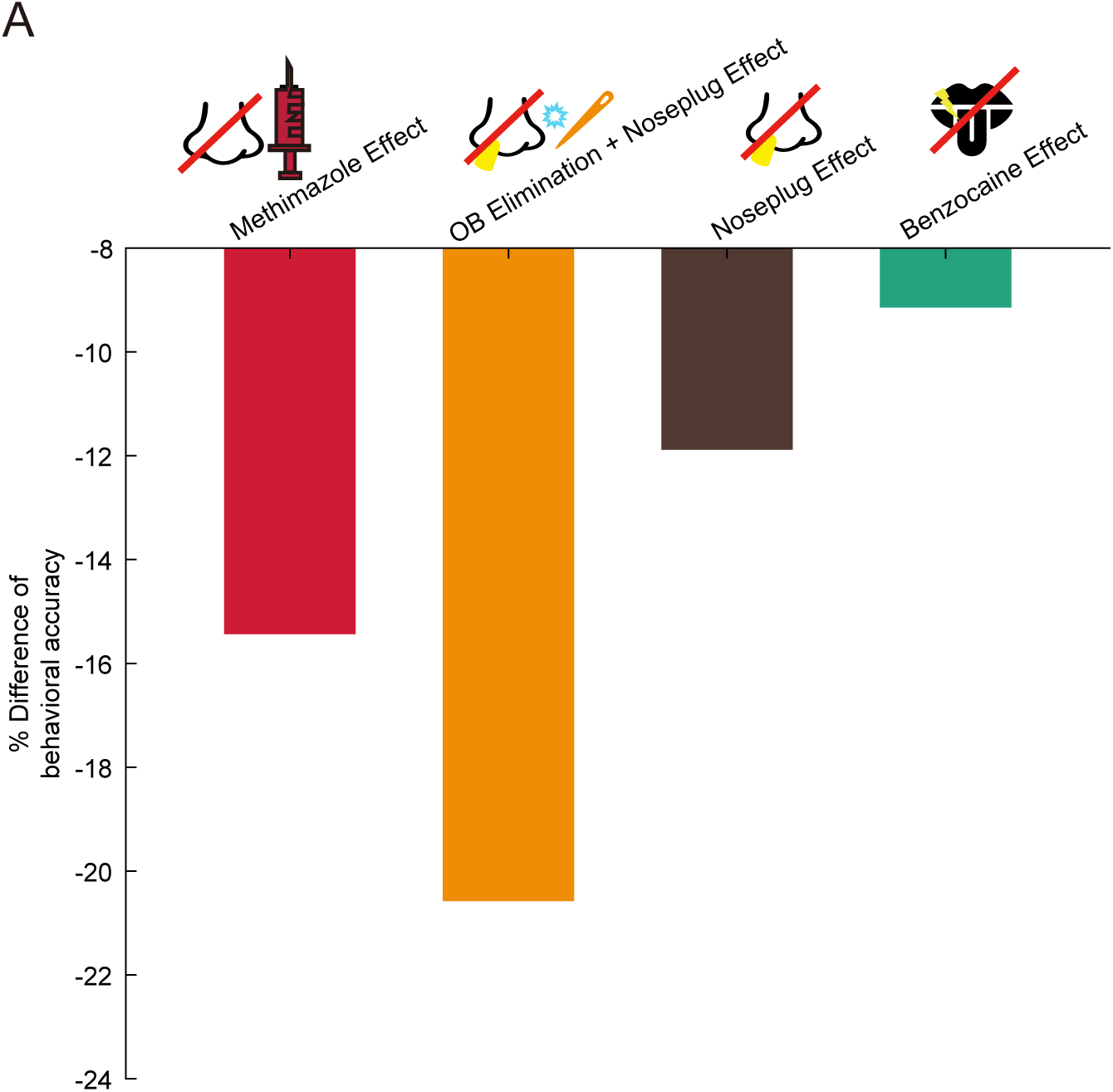
Effect of the accuracy in various validation methods for the flavor discrimination task. (A) Bar graph showing the effect of each validation method on the flavor discrimination task. The effect of methimazole was calculated as the difference in the mean percent of correct responses between the methimazole and control groups at post3 session (Figure 3). The OB elimination + nose plug effect was calculated as the difference in the mean accuracy between the OB removal and control groups in the post session (Figure 5). The unilateral nose plug effect was calculated as the difference in the mean accuracy between the pre and post sessions in the control group (Figure 5). The effect of benzocaine was calculated as the difference in mean accuracy between the pre and post sessions (Figure 6).

## Discussion

The main purpose of the study was to determine whether mice could discriminate and sense flavors. To investigate the possibility of flavor discrimination in mice, we performed a flavor discrimination task (Figure 1). The mice were able to discriminate between the presence and absence of flavors in approximately 21 sessions (Figure 2). To determine what prompted the choices made by the mice, we applied a GLM analysis to the behavioral data and showed that the mice performed the task based on the stimulus. To clarify whether this stimulus was discriminated based on olfaction, we blocked the olfactory pathway in two different ways and examined the percentage of correct responses to the flavor discrimination task (Figures 3–5). When the olfactory pathway was blocked, the mice could not discriminate between different flavors. However, when the gustatory pathway was blocked, the mice were still able to discriminate the presence of flavors (Figure 6). By blocking the olfactory system, the accuracy of this behavioral task decreased remarkably (Figure 7). These results indicate that mice can discriminate flavors depending on their olfaction and that mice are a useful animal model for elucidating the neural circuitry of the brain during flavor perception.

### Development of flavor discrimination task

Rozin (1982) hypothesized the “a dual sense modality”, which implies that the perception of odorants depends on two types of odor routes^28^. Following Rozin’s hypothesis, several flavor studies were conducted^1,6,8,17–19^. Human functional magnetic resonance imaging studies have shown that these are key regions for flavor perception in the medial prefrontal cortex (mPFC), orbitofrontal cortex (OFC), olfactory cortex (OC), insular cortex(IC) and gustatory cortex (GC) ^1, 6–9^. However, the extent to which the neural circuit mechanisms can be explored in humans is limited. This is because human neuroimaging is limited by temporal and spatial resolutions and cannot resolve neural circuit levels, making it impossible to manipulate the circuitry of the brain. To address this problem, it is necessary to conduct research using animal models. Monkeys have been used for flavor research^29, 30^, but their use has limitations owing to the number of animals available, time, and budget constraints. Conducting research on rodents is preferable to using them as model animals because various genetic techniques can be used^31^. There is supportive data that rodents perform flavor-based tasks during research^17–19^. However, there has been no detailed analysis about the basis on which rodents discriminate flavors, and it is unclear whether they are truly capable of perceiving flavor. First, a fixed-head system limits active behavior in the natural environment, which is important for flavor perception. Second, previous task protocols have not conducted sufficient experiments to test whether impairing the sense of olfaction in rodents has an adverse effect on their flavor perception. To provide evidence of olfactory discrimination, they performed OB removal, but were unable to examine the physical conditions of the rodents associated with olfactory loss^20, 21^. Third, by dissolving odor molecules in water, they could not fully rule out the possibility that rodents could discriminate based on taste or somatosensory perception.

To overcome these problems, we developed a new flavor-discrimination behavioral task for mice in a free-moving behavioral state. The free-moving behavioral state enabled the measurement of mouse reactions and movement times in a more natural state. We conducted detailed mathematical modeling to determine the stimuli that the mice discriminated against for flavor. The majority of action variance was accounted for by the stimulus (Figure 2I). Moreover, we controlled for three potential confounding factors: olfactory, taste, and somatosensory cues. We avoided external odor cues from the flavored sucrose water by always placing the same odor dissolved in flavored sucrose water around the stimulus port before presenting the flavored sucrose water and non-flavored sucrose water. To block the olfactory pathway, the mice were intraperitoneally injected with methimazole. Intraperitoneal injection of methimazole does not affect regions other than OSNs^24^. Regeneration of the OE occurs gradually over 28 days with every few cells, rather than it recovering as a whole all at once^24^. In this experiment, although the OSNs almost completely disappeared after methimazole administration, the mice were able to perform the behavioral task immediately after 28 days of recovery (Figure 4), indicating that the OE olfactory map for flavor perception was maintained after 28 days of recovery. These results suggest that, even though the wiring between the OE and OB is new, the circuit for flavor perception has already been formed by previous learning. This can be reshaped based on feedback from higher-order regions (i.e., the piriform cortex). We also used another olfactory deprivation method that combined unilateral OB removal and opposite naris-occlusion. Both methods significantly decreased the accuracy of the flavor discrimination task (Figure 5). Benzocaine was directly applied to the exposed dorsal surface of the tongue. Behavioral performance after administration was not significantly different from that before administration in the flavor discrimination task (Figure 6). These results suggest that the mice could discriminate the flavor based on odor (Figure 7), indicating that olfaction is most important for the flavor discrimination task.

In the future, this task will provide operant conditioning and recording of the neural activity of the brain based on the knowledge that mice can exactly discriminate based on flavor. Furthermore, by applying optogenetic approaches to brain regions such as the mPFC, OFC, OC, IC and GC, which are thought to be important for flavor perception, we expect to reveal the circuit mechanisms of flavor perception.

### The importance of revealing the flavor sense and flavor neural circuitry

Flavor perception has received much attention because it can be markedly altered as an early symptom of a disease and is a way to check one’s health status^32–34^. The initial symptoms of flavor loss also occur in SARS-COVID-2^33, 34^. Flavor loss remains a sequela of SARS-COVID-2 and is one of the causes of decreased quality of life^35, 36^. One reason for this is that flavor plays an important role in deliciousness of food. These findings imply that understanding the neural circuitry underlying flavor has important implications for the quality of human life.

Previous flavor studies have shown that dorsal OBs under anesthesia respond differently to the same concentration of odor in orthonasal and retronasal olfaction and that the predominant location of the glomerulus is different^37^. In addition, various regions of the OC have revealed a previously unexplored process of olfactory information processing^38–40^. Based on these findings, we speculate that there may be a specific flavor pathway in the OC that provides information on the differences in glomerular response sites in the OB depending on the odor pathway. Furthermore, studies using a cannula directly in the oral cavity revealed that the GC is involved in the memory of retronasal olfaction^41^. Flavor perception is often mistaken for taste^3^. We assume that this phenomenon is due to the fact that flavor perception is perceived as a separate sensation by integrating it in the circuit including OC-GC in the information processing of the brain. By clarifying and developing the neural circuit of flavor perception, we can achieve a subjective sense of deliciousness^3^.

### Limitations of the Study

This study had several limitations. First, we did not measure breathing patterns. Since it is very important to understand the input pattern of breathing in the flavor sense, we will perform this in a future study. Moreover, it is necessary to record neural activity to reveal the neural circuit mechanisms involved in flavor perception. In addition, optogenetic tools must be used to intervene in the perception of flavor in the brain.

In conclusion, we examined the hypothesis that mice have a sense of flavor and can discriminate it. Using a variety of mathematical and pharmacological techniques, we found that mice discriminated between test stimuli based on flavor. The development of such flavor research could potentially improve our understanding of the experience of eating and ways to shape it towards more flavorful and healthier diets^1^.

## STAR Methods

### Animals

All experiments were performed on adult male C57BL/6 mice (9 weeks old, weighing 20–25 g) purchased from Shimizu Laboratory Supplies Co., Ltd., Kyoto, Japan. The mice were individually housed in a temperature-controlled environment with a 13-hr light 11-hr dark cycle (lights on at 08:00 and off at 21:00). They were provided with water after the training and recording sessions to ensure that their body weights did not exceed 80% of their initial weight, and food was supplied ad libitum. All experiments were performed in accordance with the guidelines for animal experiments at Doshisha University, and with the approval of the Doshisha University Animal Research Committee.

### Apparatus

We used a behavioral apparatus controlled by the Bpod State Machine r0.5 (Sanworks LLC, NY, USA), an open-source control device designed for behavioral tasks^42, 43^. The apparatus consisted of a custom-designed mouse behavior box with two nose-poke ports on the front wall. The box was contained in another soundproof box (BrainScience Idea. Co., Ltd., Osaka, Japan) equipped with a ventilator fan to provide adequate air circulation and low background noise. Each of the two nose-poke ports had a white light-emitting diode (LED) and an infrared photodiode. Interruption of the infrared beam generated a transistor-transistor-logic pulse, signaling the entry of the mouse head into the port. The odor delivery port was equipped with stainless steel tubing connected to a custom-made olfactometer^44^. Amyl acetate and eugenol (Tokyo Chemical Industry Co., Ltd., Tokyo, Japan) were used as orthonasal masking odors. These odors were diluted to 10% in mineral oil and further diluted to 1:9 via airflow. Flavored sucrose water was used to dissolve 0.01% of the same orally masked odor in 10 mM sucrose water. The sucrose water used as a reward contained 40 mM sucrose. Water delivery was based on gravitational flow and was controlled by a solenoid valve connected via Tygon tubing to stainless-steel tubing. The conditioned stimulus water (6 μl) and the reward amount (12 μl) was determined by the opening duration of the solenoid valve and was regularly calibrated.

### Training procedure

All behavioral tasks were conducted in one session per day. During the initial training, the mice were habituated to the experimental system and learned about the reward ports. During the first training session, mice were habituated to the mouse behavior box. Water was provided as a reward in the stimulus port illuminated by an interior LED light. In the following session, the mice were poked into the stimulus port illuminated by an interior LED light and were presented with flavored sucrose water (6 μl). The mice were rewarded if they poked their nose into the reward port. Next, the mice were trained to move from the stimulus port to the reward port within 5s over the subsequent two to five sessions. The mice were trained to detect flavor or non-flavor, which were presented on a pseudo-randomized schedule (equal numbers within each block of 20 trials, ensuring different orders of presentation for go and no-go trials within each of the 20 trial blocks). Upon presenting flavor, the animal poked their nose into the reward port in 5 s and achieved 40 mM sucrose water as a reward (12 μl), whereas upon presenting non-flavored sucrose water, they withdrew. When the animals met the criterion of 75% choice accuracy, they were advanced to the final stage of training. This procedure lasted for five to ten sessions. Finally, the same odor dissolved in flavored sucrose water was delivered when the mice poked their nose into the stimulus port. The same odor was delivered despite the conditioned stimulus. This training lasted for 7–21 sessions to meet the above criteria.

### Flavor discrimination task

After a 5 s inter-trial interval, each trial began with the illumination of the LED light at the right odor port, which instructed the mice to poke their nose into the stimulus port. A nose poke in the stimulus port resulted in the delivery of orthonasal masking odor for about 350 ms. Next, either 6 μl of flavored sucrose water or sucrose water was delivered in the stimulus port. The mice were required to lick the conditioned stimuli. After retronasal stimulation, the LED light was turned off, and the mice could withdraw their nose from the stimulus port. If flavored sucrose water (go cue water) was present, the mice were required to move to and poke their nose into the left water-reward port within 5 s. At the reward port, the mice were required to maintain nose poking for 500 ms before the water delivery began. Then 12 μl of sucrose water was delivered as a reward. If non-flavored sucrose water (no-go cue water) was presented, mice were required to avoid entering the reward port for 5 s. The accuracy rate was calculated as the total percentage of successes in the go and no-go trials per session.

### Methimazole administration

To ablate the existing olfactory sensory neurons, the mice were intraperitoneally injected on day 0 with methimazole (75 mg/kg; Sigma-Aldrich) dissolved in saline^23^. The mice in the control group were intraperitoneally injected with saline. To compare data within a group, methimazole was administered after 3 days of saline administration to the mice in the control group. Randomly selected mice were allowed to rest on ad libitum water and food after completing the task 3 days after methimazole administration. To avoid a decrease in performance due to forgetting the behavioral task, we restricted the mice to water conservation for 20 days after methimazole administration and started the same behavioral task again on day 21.

### Unilateral olfactory elimination surgery

Adult male mice (18–22 g at the time of surgery) were anesthetized with medetomidine (0.75 mg/kg ip), midazolam (4.0 mg/kg ip), and butorphanol (5.0 mg/kg ip) prior to the olfactory elimination surgery. A rostral–caudal midline incision was made in the skin overlying the dorsal surface of the skull and a small burr hole (2 mm in diameter) was drilled in the left skull, 6 mm rostral to the bregma. The left OB was destroyed with a 10 μl pipette tip, removed by aspiration through a 16 gauge needle, and ofloxacin ointment was inserted into the cavity to control bleeding, taking care not to cause damage to other areas. A sham operation was performed in the same manner, but the bulbs were left intact. The wound was closed with Vicryl sutures, and animals were administered atipamezole (0.75 mg/kg ip) to reverse the effects of medetomidine and reduce the recovery period. Behavioral training was initiated 3 days after surgery (Figure 5A). Some mice died after the surgery to remove the OB, so they were excluded from the analysis.

### Nostril occlusion

A custom-made 10 mm silicon tube was inserted into the OB opposite the nostril of each mouse using a previously reported procedure^25, 26^ with tweezers before the 1h behavior task (Figure 5A). The silicone tube was filled with glue and ligature threads, with the threads protruding slightly from the tube. We observed the ligature threads in which the silicon tube was inserted into the right nose before the behavioral task.

### Benzocaine administration

First, the mice performed 50 task trials. Immediately after completion, 20% benzocaine (BEE BRAND MEDICO DENTAL.CO.,LTD., Osaka, Japan) was administered into the oral cavity, keeping the mice secured by hand. Subsequently, mice performed a behavioral task (Figures 6A and S3A).

### Data analyses

All data analyses were performed using the built-in software of MATLAB 2021a (MathWorks, Inc., MA, USA).

Box plot: For all box plots, the central mark is the median, the box edges are the 25th and 75th percentiles, and whiskers extend to the most extreme data points not considered as outliers (points 1.5 × interquartile range away from the 25th or 75th percentile).

Histogram: The response time histogram was calculated from histogram counts using a MATLAB function. As a parameter of detail, a bin size of 100 ms was used to calculate the histogram of relative probability.

Generalized linear models: To estimate the impact of task parameters on behavioral performance, we conducted a generalized linear model (GLM) analysis of go/no-go actions (Figure 2I and S1C). In these models, we used the identity function as the link function. For the go/no-go action GLM analysis, we prepared trials in which the participants performed peripheral choices. Task parameters included binary stimulus predictors (1 indicated the presence of flavor stimulus and 0 indicated the presence of non-flavor stimulus), response time (0 to 1 normalization), previous reward (1 indicated a reward, 0 indicated no reward), two trials before reward (1 indicated a reward, 0 indicated no reward), previous action (1 indicated the go action, 0 indicated the no-go action), two trials before action (1 indicated the go action, 0 indicated the no-go action), and thirsty state (normalized the amount of water drinking up to that point from 0 to 1). The model fit these predictors. To quantify the impact of each task parameter, we calculated the difference between the explained variances (R^2^) of the full and partial models^45^. We used 5-fold cross-validation by leaving a 20% subset of trials for prediction to avoid overfitting. This procedure was repeated 500 times. The partial model lacked the target task parameters. Behavioral data from mice with fewer than 5 trials of either go or no-go behavior were excluded from the analysis because they could not be GLM-fitted (methimazole administration group and OB removal group).

Statistical analyses: Data were analyzed using MATLAB 2021a. Statistical methods for each analysis are described above in the Results section and in the figure legends. For the pooled data (Figures 2F, 2G, 2H, 6E, 6F, S1B, S2F, S2G, S3E, and S3F), the correspondence between the before and after data was lost, and we conducted a test without correspondence. Biological replicates for the histological studies are shown in the figure legends^46^.

## Data availability

Data supporting the findings of this study are available from the corresponding author upon request.

## Code availability

The custom code used for the analyses in this study is available from the corresponding author upon request.

## Acknowledgments

We thank the Sakurai lab members for valuable discussions. K.S. was supported by JSPS KAKENHI Grant Numbers 18J21358, 21K06440, 22K15232, 23H04369, Lotte Research Promotion Grant, KEY COFFE Foundation, and Kobayashi Foundation. H.M. was supported by the Japan Science and Technology Agency (JST), PRESTO Grant Number JPMJPR21S9, JSPS KAKENHI Grant Numbers, 16K14557, 19K06963, 21H05833, 23H04368, Lotte Research Promotion Grant, the Naito Foundation, and the Takeda Science Foundation. Y.S. was supported by JSPS KAKENHI Grant Numbers 16H02061, 18H05088, 20H00109 and 20H05020.

## Author contributions

K.S. and H.M. designed the experiments, and K.S. and H.M. performed the experiments. K.S., Y.T., Y.O., K.M., J.H., and H.M. performed the data analysis. K.S., Y.T., and H.M. wrote the paper. Y.S. supported and advised the project.

## Supplementary figure legends

**Supplementary Figure S1. Details of the flavor identification task and a schematic of the encoding model fitted to the behavioral variables, related to Figure 2.**

(A) Expanded view of the stimulus port in the flavor discrimination task. The pipe of the external odor stimulus is attached to the top of that of the flavor stimulus.

(B) The boxplots of response times for Hit and CR, Miss and FA, and correct (Hit and CR) and false (Miss and FA) in the last training session, from left to right, respectively (n.s., not significant, ***p < 0.01, n.s., from left to right, respectively, Wilcoxon rank sum test).

(C) Schematic of the encoding model fitted to behavioral variables (see STAR Methods).

**Supplementary Figure S2. Changes in the response time and moving time by intraperitoneal administration of methimazole, related to Figure 3.**

(A) The time course of average behavioral accuracy after methimazole administration in the control group (gray line) (**p<0.01; ***p<0.001, Tukey’s test). Error bars show SEM.

(B) Total percentage bar showing the proportion of the behavioral action in the pre and post3 sessions of the control group (blue, Hit; light blue, Miss; red, correct rejection [CR]; light red, false alarm [FA]).

(C) Effect of task parameters on behavioral variability with methimazole administration in the saline group using a GLM fitting for go or no-go actions (*p < 0.05, two-sample t-test, see STAR Methods).

(D) Histogram of the response time in the pre session.

(E) Same as in (D) but in post3 session.

(F) The average response time of Hit, Miss, CR and FA in the pre and post3 sessions of the methimazole group (**p < 0.01, ***p < 0.001, Wilcoxon rank sum test).

(G) The average moving time of Hit and FA in the pre and post3 sessions (**p < 0.01, ***p < 0.001, Wilcoxon rank sum test).

**Supplementary Figure S3. Effects of benzocaine on the taste discrimination task, related to Figure 6.**

(A) Schematic of the schedule of the benzocaine group (n=5 mice) in the taste discrimination task. The pre session (50 trials) represents the session before the benzocaine procedure. The post session (50 trials) represents the session immediately after the pre session, when benzocaine was administered to all the mice.

(B) The average behavioral accuracy in the pre (green) and post (black) sessions of the benzocaine procedure. The behavioral accuracy in the pre session was not significant compared to the post session (*p < 0.05, paired t-test). Error bars show SEM.

(C) Percentage bar showing the percentage of the behavioral action in the pre and post sessions (blue, Hit; light blue, Miss; red, correct rejection [CR]; light red, false alarm [FA]).

(D) The average of the total number bar showing the actual number of trials in which behavioral actions were taken and corresponding to the total percentage bar (Figure S3C) (*p<0.05, paired t-test). Error bars show SEM.

(E) The boxplot of response time for individual actions in the pre (green) and post (black) sessions (*p<0.05, Wilcoxon rank sum test).

(F) The boxplot of moving time for Hit and FA in the pre (green) and post (black) sessions (***p < 0.001, Wilcoxon rank sum test).

(G) Effect of task parameters on behavioral variability with benzocaine administration using a GLM fitting for go or no-go actions. Each average Δ*R*^2^ (green, pre session; black, post session) is shown (*p<0.05, paired t-test). Error bars show SEM.

