## Supplementary figures and images for "An intra-oral flavor discrimination task in freely moving mice"

### Supplemental Figures 1-3

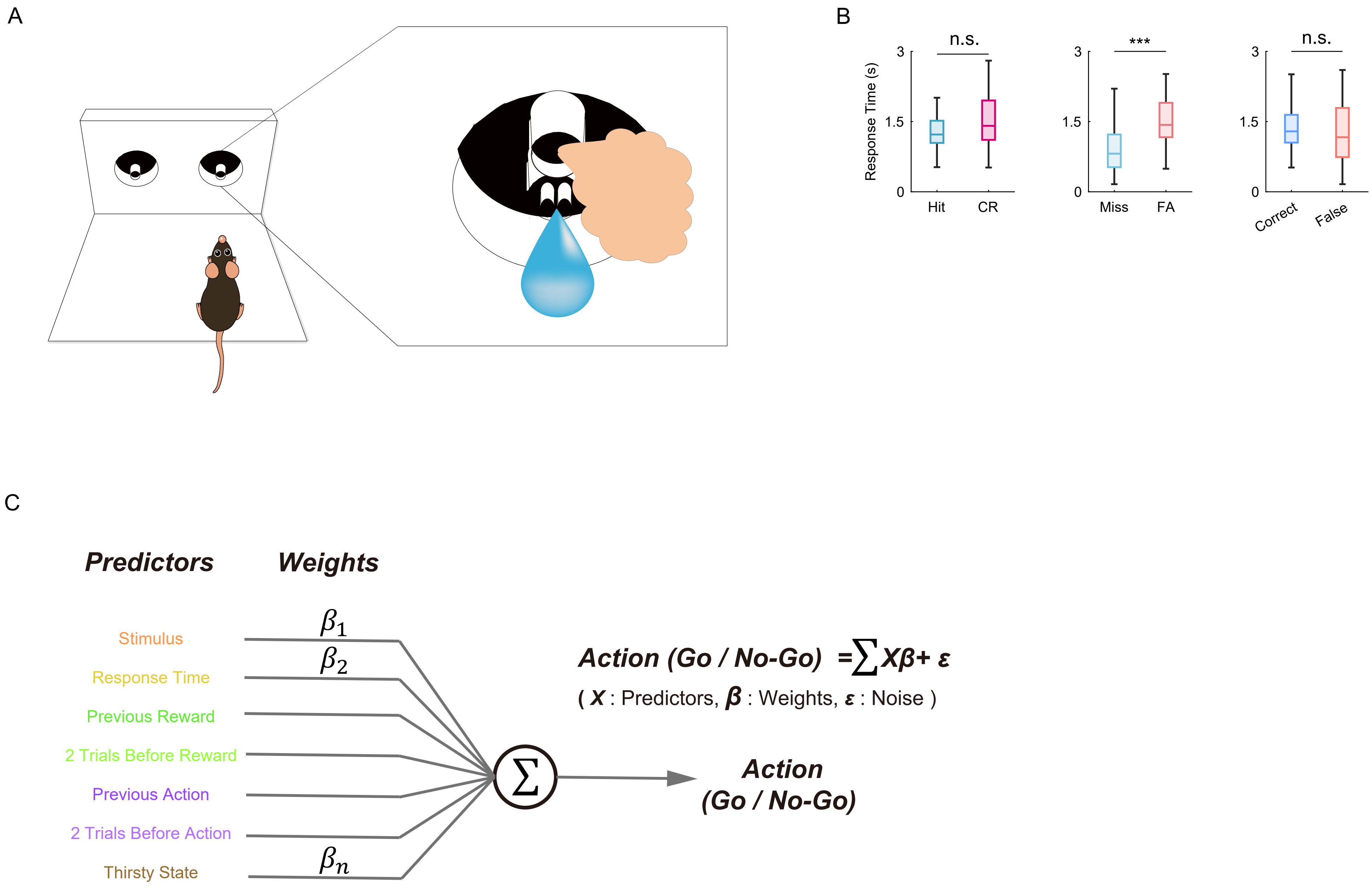

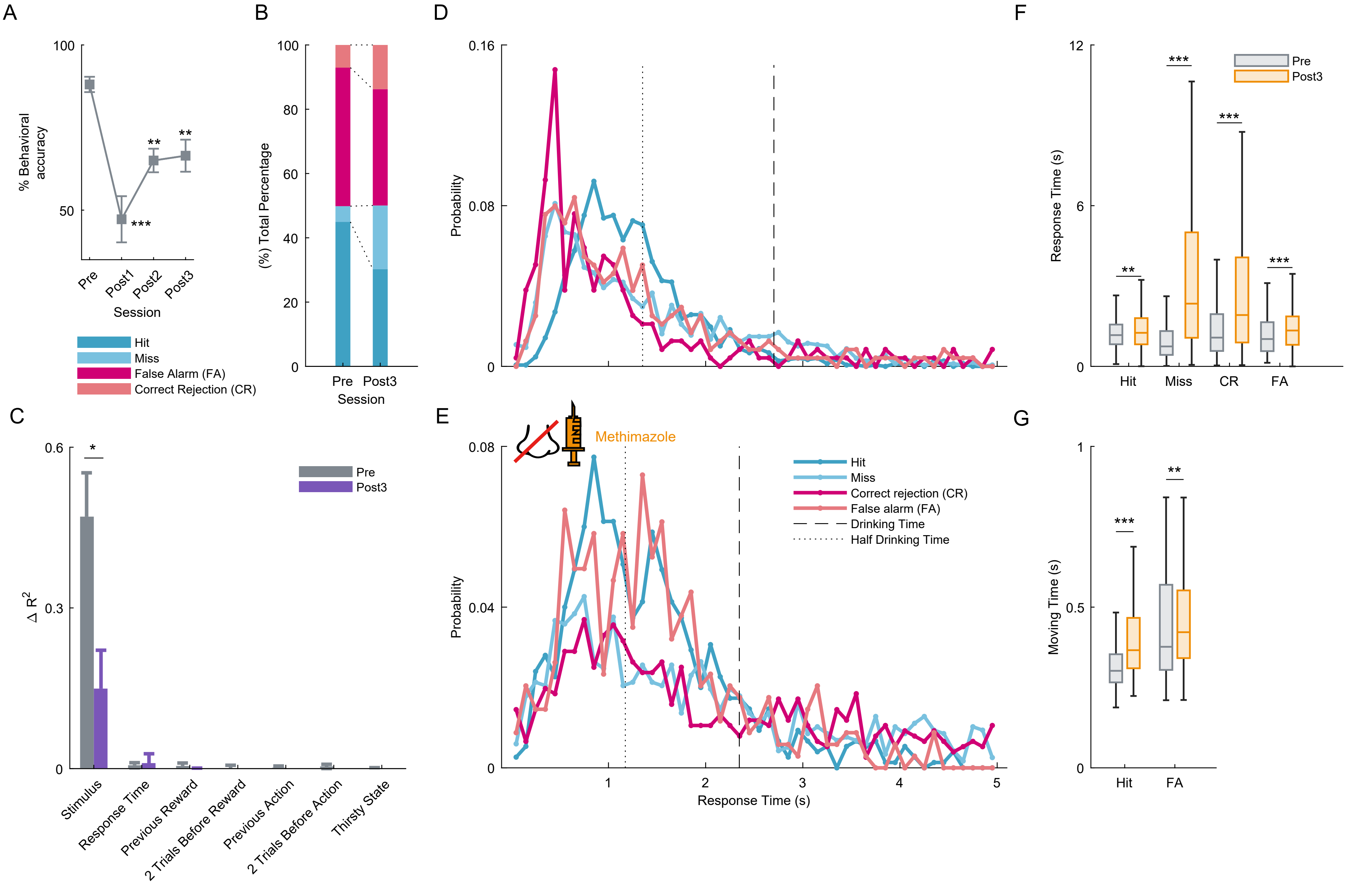

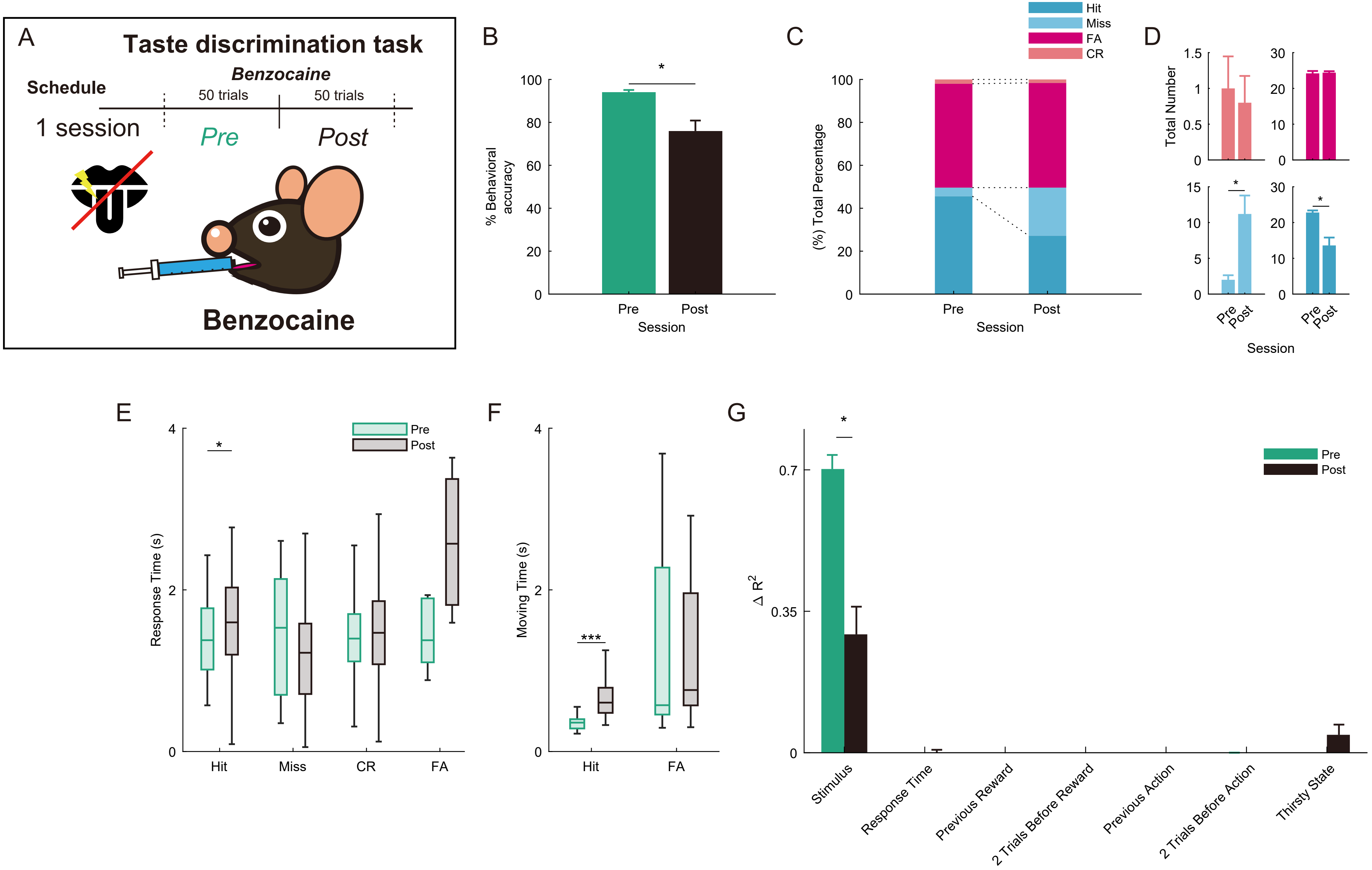
